# Xenobiotic compounds modulate cytotoxicity of Aβ amyloids and interact with neuroprotective chaperone L-PGDS

**DOI:** 10.1101/2020.01.27.920884

**Authors:** Kimberly Low Jia Yi, Margaret Phillips, Konstantin Pervushin

## Abstract

A positive association of the exposure to different classes of xenobiotics such as commonly prescribed drugs and polycyclic aromatic hydrocarbons (PAH) typically those found in air pollution-related particulate matter with Alzheimer’s disease (AD) may point to direct physical interaction of those compounds with the amyloid formation and clearance processes. In this study, for the first time, we provide evidence of such interactions for three representative compounds from prescription drugs and air pollution, e.g. anticholinergic drugs Chlorpheniramine, a common antihistamine, and Trazodone, an antidepressant as well as 9,10-PQ, a common PAH anthraquinone abundantly present in diesel exhaust and associated with AD. We demonstrate that these three compounds bind to the lipophilic compound carrier and neuroprotective amyloid beta (Aβ) chaperone lipocalin-type prostaglandin D synthase (L-PGDS) with high affinity attenuating its neuroprotective chaperone function with Chlorpheniramine exhibiting markedly stronger inhibitory effects. We also show that these compounds directly interact with Aβ(1-40) increasing the fibril’s yield with altered fibril morphology and increased the cytotoxicity of the resulting fibrils. We propose that exposure to some xenobiotics in the peripheral tissues such as gut and lungs might result in the accumulation of these compounds in the brain facilitated by the carrier function of L-PGDS. This might lead to attenuation of its neuroprotective function and direct modification of Aβ amyloid morphology and cytotoxicity. This hypothesis might provide a mechanistic link between exposure to xenobiotic compounds and the increased risk of Alzheimer’s disease.

**Graphical Abstract:** 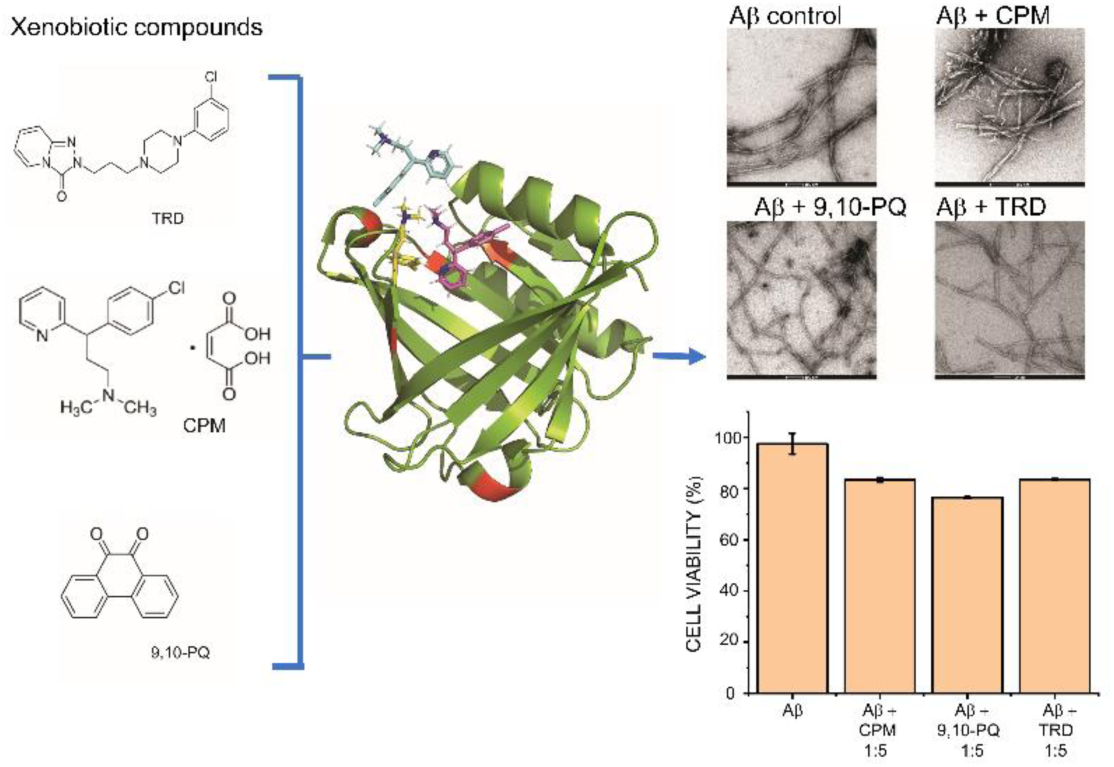

## 1. Introduction

Alzheimer’s disease (AD) is a progressive neurodegenerative disorder where symptoms such as cognitive and memory loss gradually exacerbate over the years [1]. AD together with vascular dementia is the leading cause of death among females in England (UK Office for National Statistics). The amyloid cascade hypothesis [2, 3] is one of the most popular hypotheses to explain the pathogenesis of AD. It postulates that the aggregation of Aβ peptides in the extracellular plaques is the key pathological event in AD [2]. These plaques are the result of self-assembly of Aβ fibrils which in turn are formed via the aggregation of Aβ peptides [1]. Despite general views that the accumulation of Aβ plaques in the amyloid cascade hypothesis is the result of the overproduction of Aβ peptides [4] [5] [1], recent research has found that the production rate of Aβ peptides in the cerebrospinal fluid (CSF) is similar in both control (6.8%/hr) and sporadic AD population (6.8%/hr) [6]. However, the clearance rate of Aβ in AD patients is 49% lower than the controls [6]. Hence, the decrease in clearance of Aβ peptides in the CSF may contribute to Aβ build-up in AD [7] [8]. This means that alteration or attenuation of the functionality in factors involved in the homeostasis of Aβ peptides might contribute to AD development [9].

Recently we determined the structure, dynamics and catalytic mechanism of human lipocalin-type prostaglandin D synthase (L-PGDS) [10] and characterized its Aβ chaperone function [11] proposed by *Kanekiyo et. al*. [12]. We showed that L-PGDS inhibits nucleation and propagation of Aβ fibrils and is capable of disaggregation of pre-formed Aβ amyloids in vitro as well as in AD brain extracts releasing proteins typically associated with plaques [11]. L-PGDS is the second most abundant protein in human CSF (after albumin) with approximate concentrations of 26 mg/l [13] [14]. It is also abundantly expressed in peripheral tissues [15]. This protein exhibits a strong capacity to bind to a variety of lipophilic ligands [16] and was used as a drug delivery vehicle from gut to brain [17]. The 3D structure of human L-PGDS exhibits a single eight-stranded β-barrel with a deep calyx capable of simultaneous binding of several lipophilic or amphiphilic ligands [10] and Aβ peptides with nanomolar affinity [12] [11]. This function of L-PGDS might facilitate the accumulation of some xenobiotics in the brain followed by peripheral exposure in gut, nasal mucosa and lungs potentially interfering with the homeostasis of Aβ peptides associated with AD.

Association studies of prolonged exposures to some active xenobiotics available such as prescription drugs [18, 19] [20] [21] or inhaled aerosol particulate matter [22-25] have shown that some compounds might significantly increase the risk of AD-associated dementia. However, the molecular mechanisms of these associations are not known. Given the fact that L-PGDS may contribute to controlling of Aβ aggregation [12] [11] and its ability to bind and transport a variety of drug-like small molecular ligands [26] [17], we hypothesize that these compounds might interfere with the Aβ homeostasis via their interaction with L-PGDS. Furthermore, some xenobiotics transported by L-PGDS may be capable of directly interacting with the Aβ peptides and modifying their morphology and cytotoxicity [27]. The morphology of amyloid fibrils found in different AD patients is evidently different from each other [28]. This morphological difference among patients may correlate with the origin and course of the disease and cytotoxicity of the amyloids [28] [29]. We propose that these xenobiotic compounds with high affinity to L-PGDS might be transported to Aβ aggregation sites and remodel the structure of the fibrils formed in the presence of these compounds. The altered fibrils might result in higher cytotoxicity towards neurons thus exacerbating the risk of AD.

In our study, we have selected two prescription drugs from the anticholinergic family, e.g. Chlorpheniramine maleate (CPM) and Trazodone (TRD) and one air pollution-related compound, 9,10-PQ. Chlorpheniramine maleate (CPM) (Fig. S1B) is a common over-the-counter, first-generation antihistamine drug. CPM tablets are used clinically for the treatment of allergic diseases and symptoms of a cold [30]. Trazodone (TRD) (Fig. S1C) is among the top 5 most prescribed antidepressant drugs in the United States (US) [31]. Even though TRD was approved by the FDA as an antidepressant drug, it is one of the most widely prescribed off-label sleep aids in the US [32]. We have also selected 9,10-PQ (Fig. S1D), a common molecule found in the PM_2.5_ category of air pollutants and one of the main components of diesel exhaust. 9,10-PQ is a type of anthraquinone that can undergo redox cycling to produce reactive oxygen species (ROS) [33]. We report the effects of these xenobiotic compounds on the neuroprotective function of L-PGDS and the direct interaction between these compounds and Aβ peptides resulting in altered amyloid morphology and increased cytotoxicity. In this study, tryptophan fluorescence quenching assays and Isothermal Titration Calorimetry (ITC) are carried out to characterize binding of the selected compounds to human L-PGDS. Thioflavin T (ThT) assays are performed to demonstrate inhibition of nucleation and fibril disaggregase functions of L-PGDS. Binding sites of the ligands to L-PGDS are mapped using NMR titrations. The potential alteration of the fibril morphology and biological activity induced by the compounds are studied by using ThT assays, Transmission Electron Microscopy (TEM) and MTT cell-based assays.

## 2. Materials and methods

Trazodone (TRD) (analytical grade) and 9, 10 – PQ (analytical grade) were purchased from Sigma Aldrich (Singapore). Chlorpheniramine, available as an OTC drug, was purchased from local pharmacy store. Synthetic amyloid β (1-40) peptide was custom ordered from ChinaPeptides (China).

### 2.1 Expression and purification of wild type, human unlabeled L-PGDS

Glycerol stock of Rosetta 2 DE3, *E.coli* cells (Novagen) with pNIC-CH vectors was prepared via transformation. The rest of the protocol was followed from the paper [10]. The concentrated protein was injected into the AKTA purifier Fast Performance Liquid Chromatography (FPLC) (GE Healthcare, USA) and further purified using Superdex 75 column in 50 mM sodium phosphate buffer (pH 7.0). Sodium dodecyl sulphate-polyacrylamide gel electrophoresis (SDS-PAGE) was run to check the purity of the different fractions obtained after the FPLC run.

### 2.2 Expression and purification of wild type, human ^15^N labeled L-PGDS

The glycerol stock of the *E.coli* cells used for the ^15^N labeled L-PGDS was the same as used for unlabeled L-PGDS. Starter culture was prepared in TB broth with kanamycin (50mg/ml) and chloramphenicol (25mg/ml) at 1:1000 dilutions and inoculated with the glycerol stock containing the transformed cells. Expression, extraction and purification of the ^15^N labeled L-PGDS was done in the same manner as the unlabeled L-PGDS.

### 2.3 Tryptophan fluorescence quenching assays

CPM, TRD and 9, 10 – PQ were dissolved in dimethyl sulfoxide (DMSO) to give a 2.0 mM and a 20 mM stock solution respectively. Various concentrations of each compound were added to L-PGDS in 50 mM sodium phosphate (pH 7.0) buffer and the final concentration of the protein was adjusted to 2 µM. Maximum volume of DMSO in each 100 µL well was restricted to 2 µL (2%). The L-PGDS-compound complex was formed by incubating L-PGDS with the respective compounds at 25°C for 1 hour. 100 µL of each complex was added to the Corning Costar® 96-well black, opaque, flat bottom plate. The intrinsic tryptophan fluorescence of L-PGDS was measured using Cytation 5 cell imaging multi-mode (Biotek Instrument Inc., USA) reader with λ_ex_=295 nm and λ_em_=340 nm. Duplicate measurements were performed

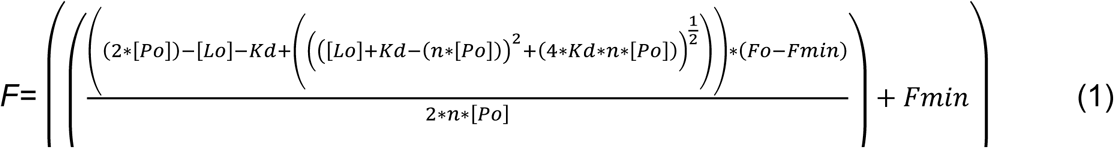

for each concentration of the three compounds. Mass law equation was used to calculate the apparent dissociation constant (K_d_) values for the drug binding to L-PGDS using the following fitting equation (eqn 1):

### 2.4 Isothermal Titration Calorimetry (ITC)

Calorimetric experiments were performed with Microcal ITC200 (Malvern, USA), in 50 mM sodium phosphate buffer (pH 7.0) at 25°C. L-PGDS (200 µM) in the injection syringe was reverse titrated into 2 µM TRD, 2 µM CPM and 2 µM 9, 10 – PQ in the cell. Titration experiments consisted of 19 injections spaced at 360 s intervals. The injection volume was 2 µl and the cell was continuously stirred at 750 rpm. The observed enthalpy changes (Δ*H*°) for binding and the dissocation constant (*K*_*d*_) were directly calculated and evaluated from integrated heats using one set of independent binding sites model available in MicroCal Origin 7.0 software [34].

### 2.5 ThT inhibition assay

ThT dye (Sigma-Aldrich, Co) was dissolved in Mili Q water and filtered through a 0.2 µm syringe filter to obtain a stock concentration of 2.3 mM. The ThT dye was diluted to give a final working concentration of 200 µM. Aggregate-free Aβ(1-40) solution at a stock concentration of 250 µM was prepared by dissolving the synthetic Aβ(1-40) powder (ChinaPeptides Co., Ltd) in 1,1,1,3,3,3-Hexafluoro-2-propanol (HFIP). The HFIP in the mixture was left to evaporate overnight to obtain the dry powder. 20uL of 100 mM NaOH was added to the HFIP treated monomeric Aβ peptide powder to further dissolve the Aβ(1-40) and remove any pre-formed aggregates. After centrifuging at 14000 rpm for 5 minutes, the supernatant was collected and 50 mM of sodium phosphate buffer was added to obtain a final working concentration of 25-50 µM monomeric solution of Aβ(1-40) peptide. The fluorescence intensity of ThT was measured using Cytation 5 cell imaging multi-mode (Biotek Instrument Inc., USA) reader with λ_ex_=430 nm and λ_em_=480 nm. The kinetics experiment was run for 72 hours at 37°C to monitor the formation of the Aβ(1-40) fibrils from the monomeric Aβ(1-40) peptide solution.

### 2.6 ThT Disaggregase Assay

Monomeric 200 µM, Aβ(1-40) peptide was grown at 37°C shaking at 200 rpm for 72 hours in an incubator to form the fibrils. For the control, 5 µM ThT dye was added to each well (100 uL) containing 10 µM preformed Aβ(1-40) fibrils. L-PGDS/drug complex was incubated for 30 minutes before adding to the wells. Duplicate measurements were performed for each condition on TECAN infinite M200 Pro microplate reader (Tecan Trading AG, Switzerland) with orbital shaking before each measurement. The kinetics experiment was run for 2 hours at 37°C to monitor the breakdown of the preformed Aβ(1-40) fibrils via L-PGDS and the L-PGDS/drugs complexes. The percentage disaggregation of preformed Aβ(1-40) fibrils was calculated using the formula:

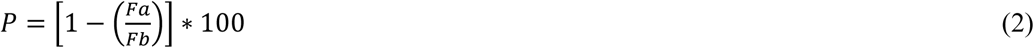

Where P is the percentage of disaggregation, *Fa* is the ThT fluorescence intensities of 10µM preformed Aβ(1-40) fibrils and *Fb* is the ThT fluorescence intensities in the presence of L-PGDS or L-PGDS-CPM complex respectively.

### 2.7 ^15^N labeled Nuclear Magnetic Resonance (NMR) titration

NMR titration experiments of L-PGDS with the CPM were acquired on Bruker Avance 700 MHz with triple resonance z-axis gradient cryoprobe at 298 K. All NMR samples contained 300 µM uniformly ^15^N-labelled L-PGDS (90%/10% D_2_O) in 50 mM sodium phosphate buffer (pH 7.0). Proton chemical shifts were referenced internally to 4, 4-dimethyl-4-silapentane-1-sulfonic acid (DSS) at 0.00 parts per million (ppm) with heteronuclei referenced by relative gyromagnetic ratios. ^1^H-^15^N HSQC spectra of ^15^N-labelled L-PGDS (300µM) was recorded in the absence and presence of drugs in a molar ratio of 1:1 and 1:4 (Protein: Ligand). These spectra were overlapped with the reference spectrum of L-PGDS alone to identify any chemical shift perturbations. Free induction decays were transformed and processed using Topspin (Bruker Biospin, USA). Assignment of the backbone nuclei of ^15^N labelled L-PGDS was carefully transferred from previously assigned spectra [10] and chemical shifts of the cross peaks were analyzed using Computer aided resonance assignment (CARA) (www.nmr.ch) [43]. The chemical shift perturbation of the affected residues was calculated using the formula:

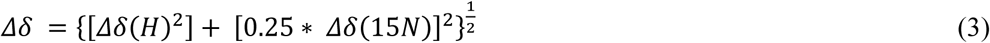

And the resulting *Δδ* was mapped onto the previously published L-PGDS crystal structure (PDB code: 4imn) to characterize binding site of the drug – CPM onto the human WT L-PGDS.

### 2.8 Transmission Electron Microscopy (TEM)

To visualize the direct interaction of CPM, TRD and 9, 10-PQ on the Aβ(1-40) fibrils formation, Aβ(1-40) monomer was incubated at 37°C for 72 h in 50 mM sodium phosphate buffer (pH 7.0). Aβ(1-40) fibrils samples treated with and without compounds were applied on copper-rhodium 400 mesh grids with 15 nm carbon coating (thickness) (prepared in-house) followed by negative staining with 2% uranyl acetate and then air dried. The samples were then viewed under FEI T12. 120 kV Transmission electron microscope equipped with a 4K CCD camera (FEI) between 48000X to 68000x magnification under low dose conditions.

### 2.9 MTT Metabolic Assay

The toxicity effects of all the amyloid fibrils were tested on SH-SY5Y human neuroblastoma cell line using [3-(4,5-dimethylthiazol-2-yl)-2,5-diphenyltetrazolium bromide] (MTT) colorimetric assay. The cells were grown in 1:1 mix of Dulbecco’s Modified Eagle Medium (DMEM) and Ham’s F-12 growth media with 5% Fetal Bovine Serum (FBS) (Gibco) containing L-Glutamine and Phenol Red in 75 mm^3^ T-75 flasks at 37°C in 5% carbon dioxide (CO_2_) environment till they achieved 70-90% confluency. After 24 hours, approximately 2500 cells were transferred into each reaction well of 96 well transparent tissue culture plate (Corning® 96 Well TC-Treated Microplates). Once the cells were stabilized, 20 µL of samples were added in each well containing 100 µL culture media. Before the fibrils were added to the cells, they underwent a washing protocol to remove excess/unbound compounds as described previously [44]. The samples were added into the cell assay and incubated for 20 hours to test their toxic effects. After incubation, 10 µL of the stock MTT reagent (5mg/ml) was added to each well. After 4 hours of conversion into formazan product, DMSO was added to dissolve the purple crystals left in the dark for 1 hour before measurement of absorbance at 570 nm with 630 nm as a subtracted reference wavelength.

## 3. Results

### 3.1 Interactions of 9,10-PQ, TRD and CPM with L-PGDS

In order to assess binding affinity of the xenobiotic compounds to L-PGDS, we carried out tryptophan quenching and calorimetric analysis on 9,10-PQ/L-PGDS, TRD/L-PGDS and CPM/L-PGDS complexes. There are three tryptophan residues in human L-PGDS: Trp43 is located at the bottom of the L-PGDS cavity while Trp54 and Trp112 are located on the H2 helix and EF loop respectively (Fig. S1) [34]. Concentration-dependent quenching of the intrinsic tryptophan fluorescence of L-PGDS was monitored as a function of the concentration of 9,10-PQ, TRD and CPM (Fig. 1A, 1B and 1C) and the extracted binding parameters are summarized in Table 1. Due to insufficient quenching of intrinsic fluorescence (Fig. 1C), binding stoichiometry of CPM to L-PGDS was not estimated from the quenching data. Even though it is unclear as to why we observe a great difference in the tryptophan quenching efficiency of the three compounds – 9,10-PQ, TRD and CPM, we suspect different binding sites in L-PGDS resulting in variable proximity of the ligand and the tryptophan residues in the complex [34].

**Table 1:** Binding affinities and thermodynamic parameters for interaction between L-PGDS with CPM and TRD

| Compound | Binding affinities and thermodynamic parameters |  |  |  |  |  |
| --- | --- | --- | --- | --- | --- | --- |
| | n | $K_d^a$<br>($\mu\text{M}$) | $\Delta G$<br>($\text{kJ mol}^{-1}$) | $\Delta H$<br>($\text{kJ mol}^{-1}$) | $-T\Delta S$<br>($\text{kJ mol}^{-1}$) | $K_d^b$<br>($\mu\text{M}$) |
| 9,10-PQ | 4.0 | $12.3 \pm 0.6$ | 43.8 | $43.7 \pm 0.73$ | -6.0 | $10.3 \pm 1.5$ |
| TRD | 3.4 | $6.5 \pm 0.2$ | -44.5 | $-48.6 \pm 0.04$ | 4.1 | $5.3 \pm 1.0$ |
| CPM | 3.0 | $5.7 \pm 0.2$ | -42.9 | $-46.8 \pm 0.04$ | 3.9 | $4.3 \pm 1.1$ |

n: Binding stoichiometry = number of molecules bound per protein molecule
a: $K_d$ obtained by ITC
$\Delta G$ : Gibbs energy change
$\Delta H$ : Enthalpy change
$\Delta S$ : Entropy change
b: $K_d$ obtained by tryptophan fluorescence quenching of L-PGDS by binding the drugs

**Fig. 1:**
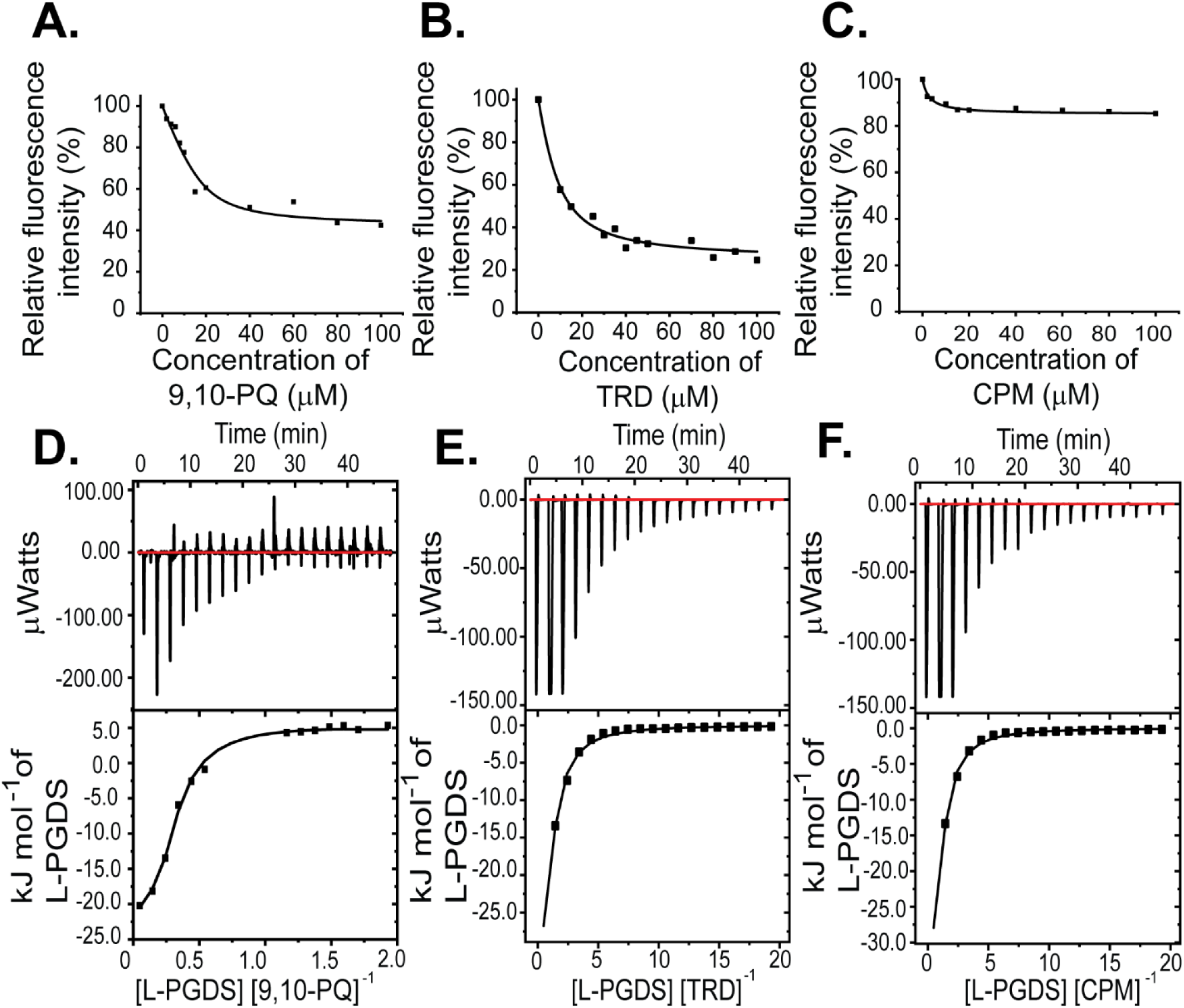
Interactions between L-PGDS and the three compounds- 9,10-PQ, TRD and CPM. Tryptophan fluorescence quenching of L-PGDS monitored at 340 nm is plotted against the various concentrations of 9,10-PQ (A), TRD (B) and CPM (C). 2 µM of L-PGDS in 50 mM sodium phosphate buffer pH 7.0 was used. Thermodynamic analysis of L-PGDS binding to 9,10-PQ (D), TRD (E) and CPM (F) in 50 mM sodium phosphate buffer pH 7.0 by ITC. L-PGDS was reversely titrated to 2 µM of 9,10-PQ, 2 µM of TRD and 2 µM of CPM. The top panes show thermograms and binding isotherms. The bottom panels show the change in heat obtained.

To further confirm binding of the three compounds to L-PGDS, thermodynamic analysis between L-PGDS and 9,10-PQ/TRD/CPM at pH 7.0 and 25°C were examined using ITC. From the negative peaks of the titration curves obtained from ITC, binding of L-PGDS to all compounds was found to be an exothermic reaction, as indicated by the favorable enthalpy changes (Fig. 1D, 1E and 1F upper panel). Integration of each peak area of the curve was plotted against the molar ratio ([L-PGDS] [COMPOUND]^-1^) (Fig. 1D, 1E and 1F lower panel). The K_d_ values obtained for the three compounds were calculated via fitting of the binding isotherms with the single set of identical sites binding model and are in good agreement with K_d_ obtained from the tryptophan quenching data (Table 1).

### 3.2 CPM inhibits the chaperone and disaggregase function of L-PGDS by occupying Aβ(1-40) peptide binding site

We used the Thioflavin T (ThT) fluorescence assay to monitor spontaneous aggregation of Aβ(1-40) in the presence and absence of L-PGDS in complex with the selected compounds. Aβ(1-40) aggregation exhibits a characteristic sigmoidal curve indicative of amyloid formation via primary nucleation, fibril elongation and secondary nucleation [11] (Fig. 2A). The elongation phase of the control 50 μM Aβ(1-40) peptide starts at ca 10 h after the lag phase reaching the steady phase at ca 30 h which is typical for fibril formation [35]. The presence of L-PGDS at 5 μM extends the elongation phase and decreases the final amount of Aβ(1-40) peptide aggregates to 40% when compared to the control (Fig. 2A). However, when L-PGDS is in the complex with CPM in a 1:3 ratio (binding stoichiometry as obtained by ITC), the reduced efficiency of L-PGDS to inhibit the Aβ(1-40) peptide aggregation is reflected by an increase in the ThT intensity of the L-PGDS/CPM complex when compared to inhibition of Aβ(1-40) with free L-PGDS (Fig. 2A). Thus, CPM binding to L-PGDS makes the complex less efficient in suppressing the primary aggregation of Aβ(1-40) peptide as compared to free L-PGDS possibly due to a partial overlap of the CPM and Aβ(1-40) binding sites. In contrast, binding of 9,10-PQ or TRD to L-PGDS does not result in any significant effect on L-PGDS chaperone activity (Fig. S2) possibly due to the lesser degree of interference of 9,10-PQ and TRD with Aβ(1-40) binding.

**Fig. 2:**
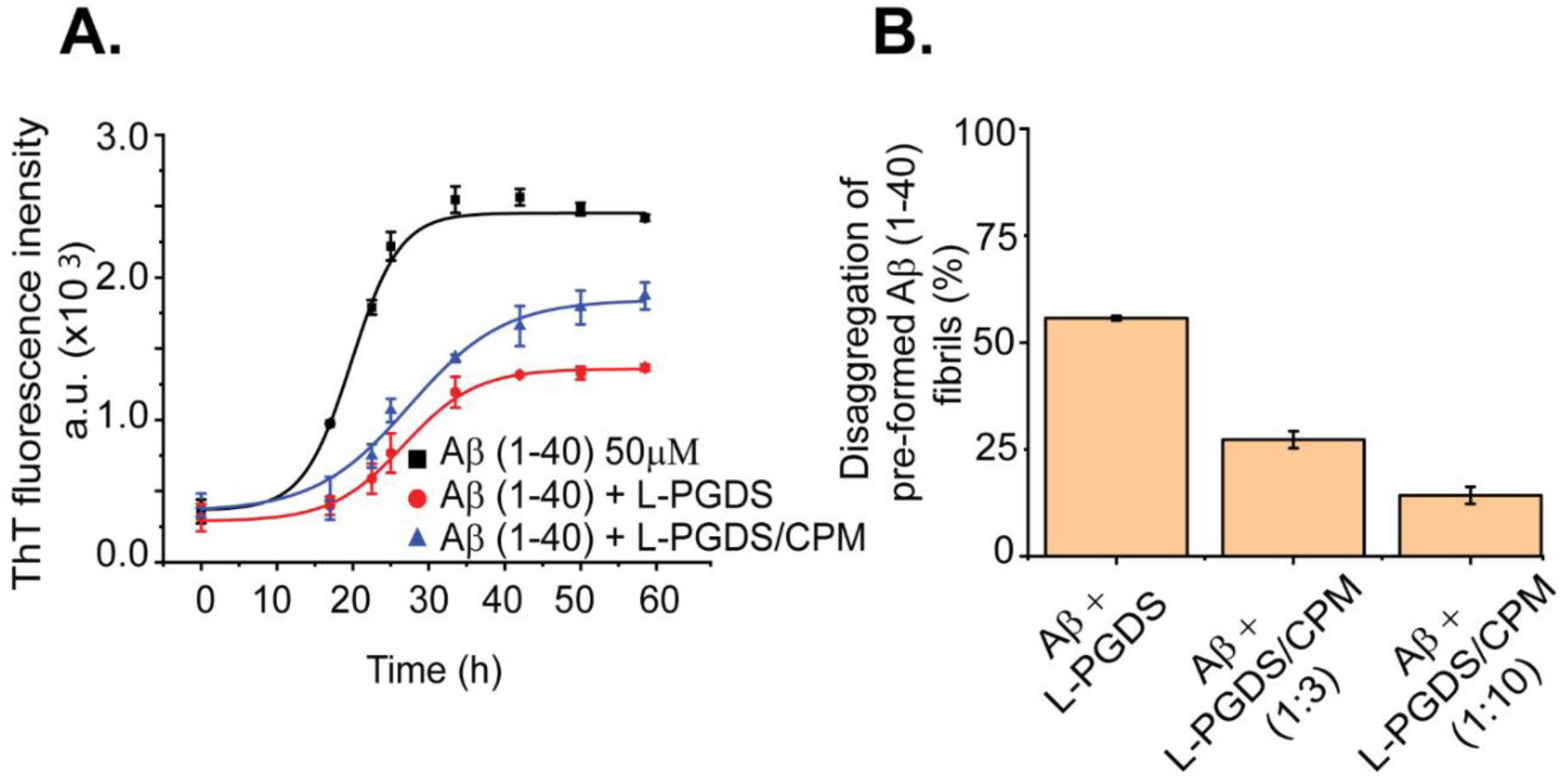
Inhibitory effects of L-PGDS and L-PGDS/drug complex on Aβ(1-40) peptide aggregation. (A) Representative time course of spontaneous Aβ(1-40) peptide aggregation in the absence (black) or presence of L-PGDS in 1:10 (protein: peptide) ratio (red) or presence of L-PGDS/CPM complex in ratio 1:3 (protein: drug) ratio (blue). (B) Disaggregation of preformed Aβ(1-40) fibrils by L-PGDS or L-PGDS/CPM complex in 1:3 (protein: drug) ratio and L-PGDS/CPM complex in 1:10 (protein: drug) ratio. Data shown are means ± SEM (n=2).

Our previous data demonstrated that human WT L-PGDS also performs disaggregase activity as part of its chaperone function by breaking down preformed Aβ(1-40) fibrils [11]. To examine if CPM binding to L-PGDS also affects its disaggregase function, we monitored disaggregation of pre-formed Aβ(1-40) fibrils with L-PGDS and (1:3) L-PGDS/CPM. The chaperone alone was able to break down up to 50% of the preformed Aβ(1-40) peptide fibrils resulting in a decrease in the ThT fluorescence intensity. At the same time, addition of the (1:3) L-PGDS/CPM complex could only break down up to ca 27% of the Aβ(1-40) peptide fibrils compared to the control (Fig. 2B). Interestingly, a higher concentration of CPM in the L-PGDS/CPM complex (1:10) exhibited higher ThT fluorescence intensity when compared to L-PGDS/CPM complex at 1:1 molar ratio. This increase in ThT fluorescence could be due to excess CPM molecules interacting with the Aβ(1-40) peptide monomers/oligomers, increasing the overall fibrillar content which competes with the breakdown of the preformed Aβ fibrils by L-PGDS thus resulting in an overall increase in fluorescence.

As only CPM displayed a significant effect on both the chaperone and disaggregase activity of L-PGDS, characterization of the CPM binding site to L-PGDS was obtained by NMR involving recording 2D ^1^H-^15^N HSQC spectra of ^15^N labeled L-PGDS with and without CPM at varying molar ratio. The assignment for the reference spectrum was carefully transferred from previously published data [10] recorded in the same sample conditions. The residues with average chemical shift perturbation higher than a preset threshold of 0.05 ppm (Fig. 3A) were mapped to the crystal structure of L-PGDS (PDB) code: 4IMN) [10] to define the interaction site of CPM with L-PGDS. Residues M64, A72, F83, E90, L131, Y132, G135, A169, T183 and E189 showed moderate chemical shift perturbations (0.05 < Δδ < 0.07) and are highlighted in orange. Residues D37, A49, G140, R144 and T152 showed large chemical shift perturbations (Δδ > 0.07) and are highlighted in red. Overall, CPM binding to L-PGDS results in significant chemical shift perturbations mainly originating from the residues located at the entrance of the β-barrel cavity of L-PGDS (Fig. 3B). The tryptophan fluorescence quenching results also point to similar binding sites for CPM positioned away from W43.

**Fig. 3:**
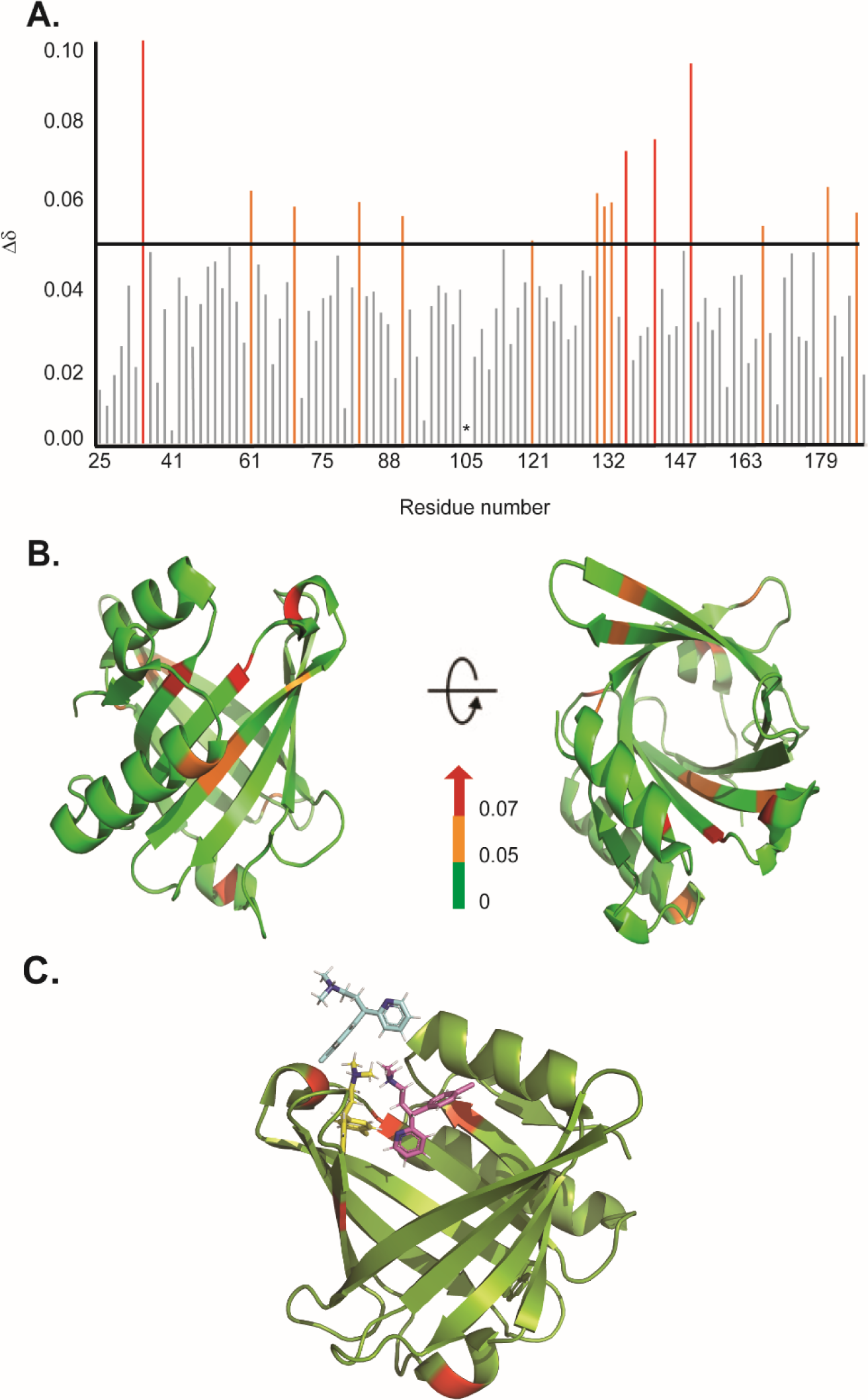
Binding sites characterization of CPM on L-PGDS, as determined by NMR titrations. (A) The chemical shift perturbations (Δδ) of backbone and amide groups of residues in L-PGDS induced by binding to CPM at a molar ratio of 1:4 (protein: drug). (B) Mapping of chemical shift perturbations (Δδ) on L-PGDS crystal structure (PDB code: 4IMN) in the presence of CPM (1:4) (protein: ligand). Red and orange show Δδ > 0.07 and 0.05 < Δδ < 0.07, respectively. In the panels, left and right images are the front and top views of L-PGDS (PDB code: 4IMN) in solution, respectively. (C) Model of three CPM molecules (blue, yellow and magenta) docking onto L-PGDS (green) obtained from online protein-ligand docking platform, HADDOCK [36]. (Color should be used)

The residues with significant CSPs (e.g. D37, A49, G140, R144) identified from our NMR titration analysis were used to guide the docking of CPM molecules in the crystal structure of L-PGDS (PDB: 4IMN) using HADDOCK, an online protein-ligand docking platform [36]. Our docking model revealed that CPM molecules bind near the β-calyx entrance of L-PGDS as shown in Fig. 3C. We discovered that the entrance of the calyx of L-PGDS can accommodate up to three CPM molecules, thus supporting the stoichiometric binding ratio estimated from our ITC measurements.

### 3.3 Xenobiotic compounds alter the Aβ(1-40) fibril morphology and increase its cytotoxicity

To further investigate the direct interaction between the selected compounds and the Aβ(1-40) we performed the ThT assay (Fig. 4A). At 1:1 stoichiometry, all three compounds induce faster fibril formation with the overall higher fibrillar content as compared to Aβ(1-40) control (Fig. 4A). The elongation phase of Aβ(1-40) fibrils start at ∼20 h, ∼25 h and ∼15 h when incubated with 9,10-PQ, TRD and CPM respectively, as compared to ∼30 h for Aβ(1-40) control. TRD, CPM and 9,10-PQ shorten the lag phase while significantly increasing the total fibril content (Fig. 4A). The increase in the fibril content was also visualized in terms of increased Aβ fibril numbers as observed by TEM (Fig. 4B-E).

**Fig. 4:**
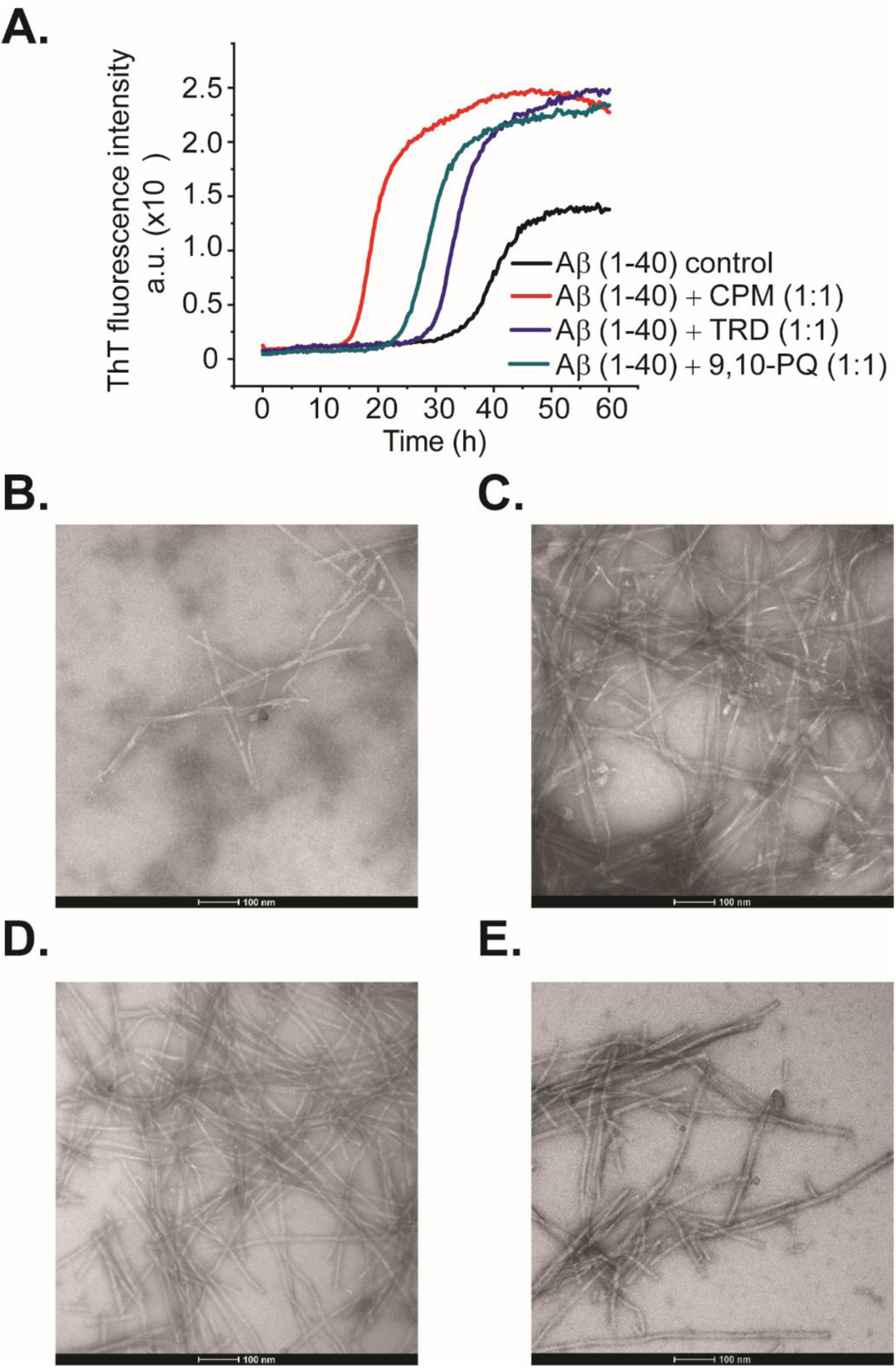
Direct interaction between CPM, TRD and 9,10-PQ with Aβ(1-40) peptides in a 1:1 ratio. (A) Representative time course of spontaneous Aβ(1-40) peptide aggregation in the absence of compounds (black) or presence of CPM in a ratio of 1:1 (red) or presence of TRD in a ratio of 1:1 (blue) or presence of 9,10 PQ in ratio of 1:1 (green). (B) TEM image of Aβ control grown for 60 h at 37°C under continuous shaking. (C) TEM image of Aβ control grown in the presence of 25 µM CPM (1:1 ratio) for 60 h at 37°C under continuous shaking. (D) TEM image of Aβ control grown in the presence of 25 µM 9,10-PQ (1:1 ratio) for 60 h at 37°C under continuous shaking. (E) TEM image of Aβ control grown in the presence of 25 µM TRD (1:1 ratio) for 60 h at 37°C under continuous shaking. (Color should be used)

Upon further increase in the concentration of CPM, TRD and 9,10-PQ to 1:5 (Aβ: compound), changes in the morphology of the Aβ(1-40) peptide fibrils were observed. The electron micrographs obtained from the TEM were analyzed using Image J software to quantify the number and size of both modulated and control fibrils of Aβ(1-40) [11]. For CPM, in addition to the increase in the total fibril content (Fig. 5B and 5F), the fibrils also became shorter and thicker in terms of width (Fig. 5B and 5E) when compared to the control fibrils. For 9,10-PQ, the total fibrillary content increases significantly (Fig. 5C and 5F) with increasing concentration of the compound along with a significant increase in the fibril length (Fig. 5C and 5D) resembling a ‘ribbon-like’ morphology (Fig. 5C). The fibrils grown in the presence of TRD do not show visible differences in number and morphology when compared to the control (Fig. 5D). Since the modified fibrils have different morphology compared to the Aβ(1-40) control, we speculate that the difference in morphology may show different biological activities of the resulting AB fibrils as it was demonstrated in the subsequent cell viability assays.

**Fig. 5:**
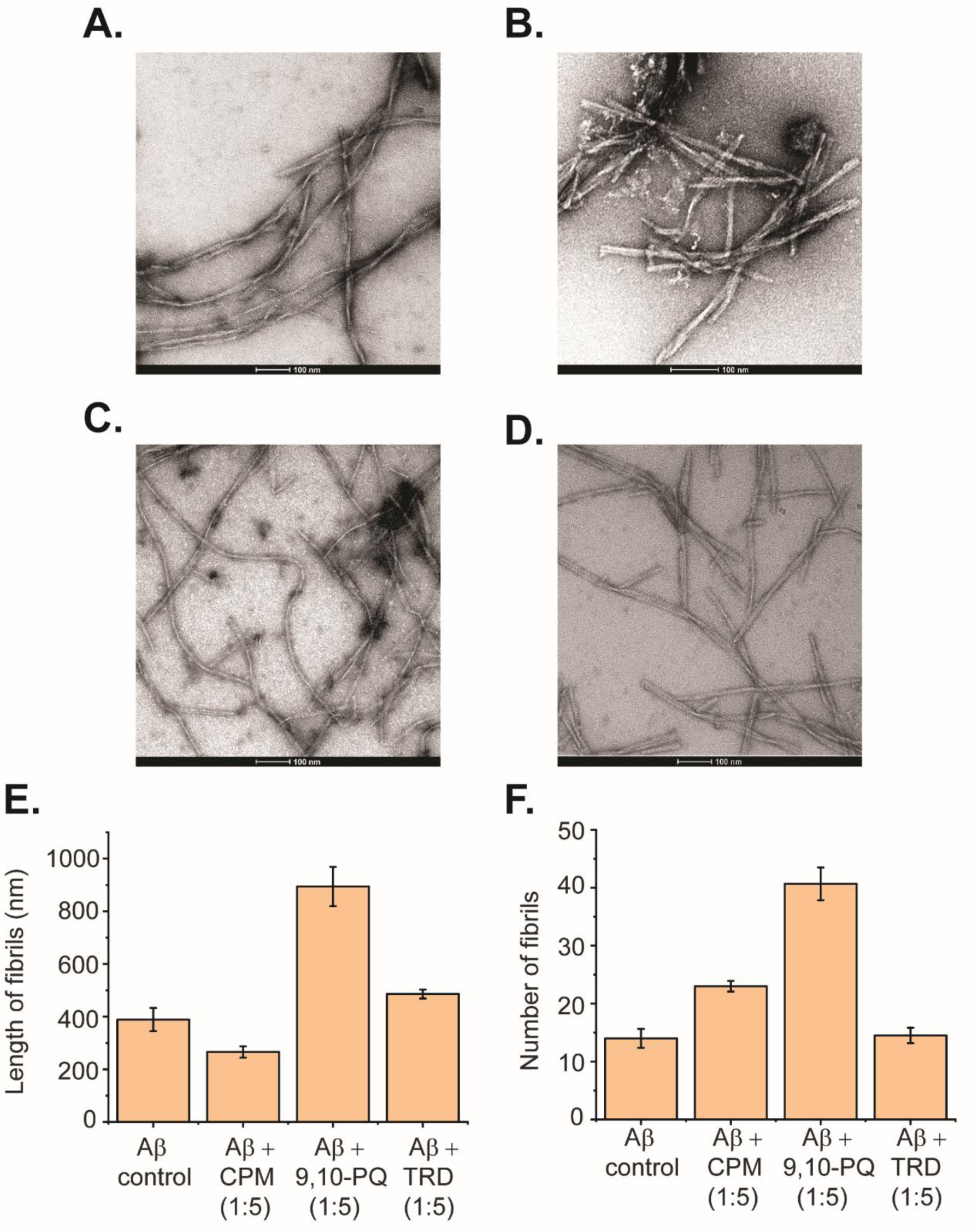
Direct interaction between 9,10-PQ and CPM with Aβ(1-40) peptides in a 1:5 ratio. (A) TEM image of Aβ control grown for 60 h at 37°C under continuous shaking. (B) TEM image of 50 µM Aβ control grown in the presence of 250 µM CPM (1:5 ratio) for 60 h at 37°C under continuous shaking. (C) TEM image of Aβ control grown in the presence of 250 µM 9,10-PQ (1:5 ratio) for 60 h at 37°C under continuous shaking. (D) TEM image of Aβ control grown in the presence of 250 µM TRD (1:5 ratio) for 60 h at 37°C under continuous shaking. (E) Length of fibrils for each condition measured from the TEM images. (F) Number of fibrils for each condition measured from the TEM images.

In addition to changes in morphology, the altered Aβ(1-40) fibrils also exhibited different effects on the disaggregase function of L-PGDS. From the ThT results it can be seen that the fibrils formed in the presence of the xenobiotic compounds display different activity as compared to the Aβ(1-40) control fibrils, where they interfere with the disaggregation efficiency of L-PGDS as shown in Fig. 6B-C. The fibrils grown in the presence of 9,10-PQ showed the greatest resistance against the disaggregase action by L-PGDS where ∼87% of the fibrils remain intact after incubation with L-PGDS. Fibrils grown in the presence of TRD (∼76%) and CPM (∼66%) also showed reduced susceptibility to L-PGDS disaggregase activity albeit to a lesser extent (Fig. 6C).

**Fig. 6.**
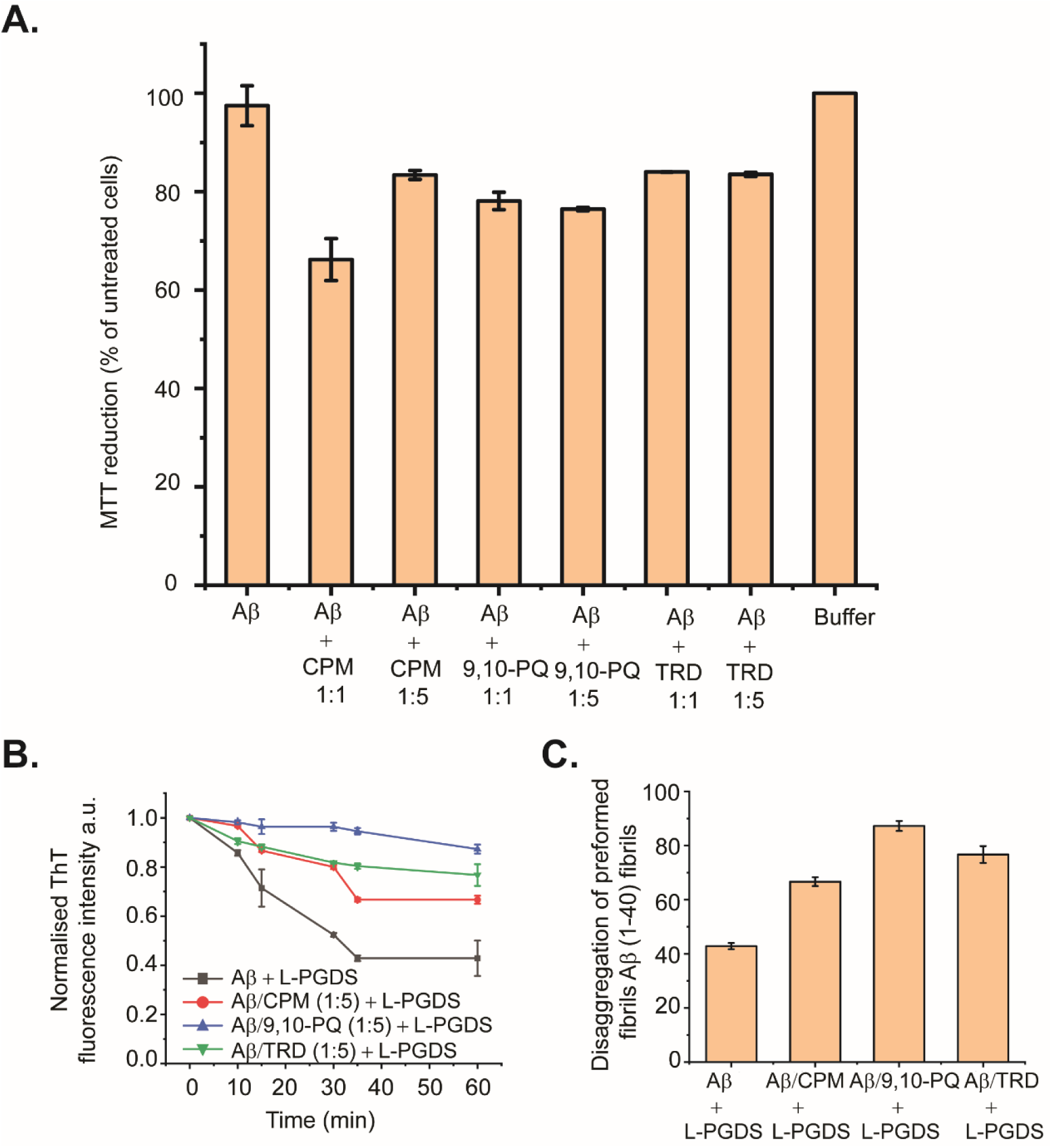
Toxicity and Biological activities of modulated fibrils (A) Various effect on the metabolic activity of SH-SY5Y cells after addition of modulated fibrils. Fibrils were grown in the presence of compounds for 72 h at 37°C and added into the cell culture media. MTT reduction was normalized to control cells. Values represented means ± SEM. (B) Representative time course of disaggregation of fibrils grown in different conditions. Control Aβ fibrils with the addition of 5 µM of L-PGDS (black), Aβ fibrils grown in 250 µM of CPM with the addition of 5 µM of L-PGDS (red), Aβ fibrils grown in 250 µM of 9,10-PQ with the addition of 5 µM of L-PGDS (blue) and Aβ fibrils grown in 250 µM of TRD with the addition of 5 µM of L-PGDS (green). (C) Disaggregation of preformed Aβ(1-40) fibrils by L-PGDS, Aβ fibrils grown in 250 µM of 9,10-PQ (1:5 peptide: compound ratio), Aβ fibrils grown in 250 µM of CPM (1:5 peptide: drug ratio) and Aβ fibrils grown in 250 µM of TRD (1:5 peptide: drug ratio). Data shown are means ± SEM (n=2).

The MTT metabolic assay was employed to investigate the cytotoxicity of the Aβ(1-40) fibrils grown in the presence of the xenobiotic compounds - 9,10 PQ, CPM and TRD in SH-SY5Y cells. The Aβ(1-40) fibril control did not exhibit any significant effect on the viability of the SH-SY5Y cells as also shown previously by Krishtal *et al* [37]. In contrast, exposure to Aβ(1-40) fibrils grown in the presence of CPM in a molar ratio of 1:1, resulted in a significant decrease in the viability of SH-SY5Y cells, a 66.19% of the control. Interestingly, the Aβ(1-40) fibrils grown in the presence of higher CPM concentration (molar ratio of 1:5 for LPGDS: CPM) only induced a slight decrease in cell viability (83.39% of the control). Exposure to Aβ(1-40) fibrils grown in the presence of 9,10-PQ in a ratio of 1:1 and 1:5 reduced the survival of SH-SY5Y cells to 78.12% and 76.46% of the control, respectively. For Aβ(1-40) fibrils grown in the presence of TRD in a ratio of 1:1 and 1:5, a drop-in cell viability to ∼ 83% of the control was observed (Fig. 6A). Overall, the MTT assays indicate that the altered Aβ(1-40) fibrils exhibit various toxicity levels when compared to the untreated Aβ(1-40) fibril control most likely due to the different morphology of the affected fibrils.

## 4. Discussion

L-PGDS exhibits multiple functions in the organism. It can act as a PGD_2_-synthesising enzyme and an extracellular transporter for lipophilic compounds [16]. Due to its ability to bind and carry lipophilic compounds, it was proposed as an efficient drug delivery vehicle to the brain dramatically increasing in-situ drugs’ availability [17]. Exposure of the peripheral tissues such as gut, nasal mucosa and lungs to the lipophilic xenobiotics with potentially strong biological activities might result in binding of these compounds to the abundant lipophilic carriers with subsequent transfer and accumulation of those compounds to the remote tissues. L-PGDS is standing out from the other lipophilic carriers by its abundance and unique ability to release its cargo into the blood [17]. For example, studies show that some air-pollution related compounds can accumulate in the brain at a concentration of 0.1-30 μM [33]. The accumulation of the biologically active xenobiotics at these elevated concentrations might interfere with a variety of processes including the neuroprotective function of L-PGDS itself and direct interference with the amyloidogenesis in the context of AD [27]. In this study, we selected three representative compounds and explored their abilities to bind to L-PGDS, interfere with its amyloid nucleation and propagation inhibition as well as its amyloid disaggregase functions. Also, we show that at elevated concentrations, which are still within the physiological range found in the brain tissues, these compounds are capable of altering the morphology of the resulting Aβ fibrils typically increasing their resistance to disaggregation and cytotoxicity.

Recently, it has been reported that the morphology of amyloid fibrils found in different AD patients are evidently different from each other [28]. It has been suggested that the morphological difference among patients correlate with the difference in disease progression [28] [29]. However, the exact reason for the different fibril morphology observed remains unknown. Here, from our study, we suggest that the modulation of fibril morphology by xenobiotic compounds could be one of the possible causes of the morphology differences observed among different patients. Given our constant exposure to these xenobiotic compounds in terms of air pollutants and common drugs, it seems reasonable to suggest that the Aβ fibrils found in AD patients can be remodeled by these compounds. Hence, exposure to these compounds might result in the different morphology of fibrils observed among different AD patients [28]. The modulated fibrils have different fibrillary morphologies and are more cytotoxic than the unmodified fibrils, indicating that certain amyloid morphologies are more toxic than others. We believe that future therapeutic interventions must take into consideration this possible morphological alteration in amyloid fibrils potentially induced by drugs. Reported here is our pilot study pointing to the direct mechanistic link between exposure to xenobiotics and change in the homeostasis of Ab peptides typically associated with AD.

We addressed the lipophilic carrier function of L-PGDS by investigating direct binding of the selected compounds to the protein. We selected compounds which have been associated with AD and belong to commonly prescribed drugs group and air-pollution-related compounds. The intrinsic tryptophan fluorescence quenching data and the ITC data establish binding of the three compounds 9,10-PQ, TRD and CPM to human L-PGDS with binding affinities of Kd ∼ 10 μM and ∼5 μM respectively (Table 1). Interestingly, we observed that CPM is the only compound that interferes with the interaction between the Aβ(1-40) peptide and L-PGDS (Fig. 2A and S2) most likely inhibiting binding of Aβ(1-40) monomers to L-PGDS. While interacting with Aβ (1-40), the chemical shifts of residues D37 and R144 of L-PGDS were significantly shifted and chemical shifts of A49 and G140 were completely attenuated [11]. The MD simulation model obtained from the study showed that Aβ(1-40) peptide interacts with L-PGDS at the entrance of the L-PGDS calyx (Fig. S3). In this study, WT L-PGDS in complex with CPM showed large chemical shifts for similar residues (Fig. 3A) and our docking model revealed CPM molecule binding to the entrance of the L-PGDS calyx (Fig. 3C) similar to Aβ(1-40). Thus, we propose that CPM exerts its inhibitory effect on the chaperone activity of WT L-PGDS by either partially or fully occupying the Aβ(1-40) peptide binding site on L-PGDS. It is also possible that CPM binding to different regions of L-PGDS results in a conformational change within L-PGDS (as shown in the large chemical shifts observed in the bottom of the calyx (Fig. 3B)) leading to an allosteric inhibition of L-PGDS’s ability to bind Aβ(1-40) monomers, oligomers or even fibrils. Our ThT results further confirm the inhibitory effect of the compound – CPM on the L-PGDS ability to reduce Aβ(1-40) aggregation as well as L-PGDS ability to break down preformed Aβ(1-40) fibrils in vitro. Further studies necessitate the need to investigate the detailed mechanism that involves the inhibitory effect of CPM on L-PGDS function. This might provide further insights into how other similar xenobiotic compounds interfere with the chaperone network within AD.

9,10-PQ, TRD and CPM were also found to directly interact with the Aβ peptide as demonstrated by the sharp increase in ThT fluorescence assays (Fig. 4A). The resulting fibrils have higher fibrillar content and longer fibrils as compared to the control (Fig. 4B-E). We propose that at a lower concentration of the selected compounds, these compounds interact with the Aβ(1-40) peptide and affect the nucleation mechanism of the Aβ peptide. The shortening of the lag phase of the ThT curve in the presence of these compounds (Fig. 4A) supports our claim that the compounds were accelerating the nucleation process of Aβ peptides as nucleation normally occurs at the lag phase of the aggregation curve for Aβ fibrils [38]. As Aβ fibrils grow via the addition of monomers at fibrils’ end [39], it is likely that these compounds interact with the short axis of the fibril and cause the elongation of the fibrils. Hence, the fibrils grown in the presence of the compounds had higher fibril content and the length of the fibrils was significantly longer. With regards to the change in morphology of the fibrils at higher concentrations of the compounds (1:5; Aβ(1-40) peptide: compounds), we propose that the large number of compounds present at such concentrations is sufficient to interact with the long axis of the fibrils (long side interactions). This interaction between the fibrils and compounds would affect the fibril packing resulting in the change in morphology of the fibrils [40] as observed in TEM images (Fig. 5A-D).

From the results of the MTT (Fig. 6A) and ThT assays (Fig. 6B), we conclude that different morphologies of fibrils caused by the interaction of the compounds can exhibit different toxicities and biological activities. We suggest that when the morphology of the fibrils is changed by the compounds, different sets of amino acids would be buried in the core of the fibrils. Hence, causing an alteration of the interaction sites of the modulated fibrils, leading to the fibrils displaying different toxicity and biological activity as compared to the unmodulated fibrils. This hypothesis is further supported by Petkova *et al* where they also suggested that the reason for different morphology leading to different biological activities could be due to the exposure of different amino acid side chains on the fibril surface [41]. It is therefore worth exploring with future studies on the detailed structure of the modulated fibril to unravel the relationship between fibril morphology and its respective toxicity and biological activity. Furthermore, it has been reported that the toxicity of Aβ fibrils could be related to the process in which the fibrils are grown during the initial nucleation process [42]. Since we have shown in our experiment that the compounds most likely interact with the Aβ monomers during the fibrillation process (Fig. 4A), the molecular mechanism on how these compounds interact and alter the structure of monomers and fibrils might also provide valuable insights for the different toxicity observed in this study.

Together, our studies show that xenobiotic compounds (9,10-PQ, CPM and TRD) might be linked to AD via transport with lipophilic carriers such as L-PGDS and disruption of neuroprotective function important for the Aβ homeostasis. We show that these compounds are capable of directly interacting with the amyloid fibrils and altering their morphology. This change in morphology resulted in various toxicity of the modulated fibrils and their respective resistance towards the disaggregation of L-PGDS. As a result, it is possible that these compounds indirectly disrupt the chaperone clearance of the fibrils. CPM is also capable of directly interfering with the chaperone network by inhibiting the chaperone and disaggregase function of L-PGDS. This resulted in a slower clearance of the aggregated Aβ load and further exacerbated the aggregation of the Aβ peptides. At present, we hope that our study would provide possible molecular mechanisms for the association between these xenobiotic compounds with AD.

## 5. Conclusion

In conclusion, we have shown that the xenobiotic compounds, 9,10-PQ, TRD and CPM, can be associated with AD via two different possible mechanisms. Firstly, they inhibit the chaperone protein such as L-PGDS, important for the Aβ aggregate clearance pathway by binding to the Aβ site and interfering with the chaperone activity of the protein. Secondly, they directly interact with the Aβ peptides (monomers and/or oligomers) to increase the Aβ peptide aggregation as well as altering the morphology of the modified fibrils and/or affecting both mechanisms at the same time. From our study, we hope to generate possible insights for future therapeutic intervention for AD, increase awareness of the potential long-term effects of the xenobiotic compounds and trigger further interest in finding more about other mechanisms that might link other xenobiotics compounds with increased AD risk. The morphological and toxicity differences of the modulated fibrils induced by xenobiotic compounds may also serve as a useful model for investigating the relationship between amyloid fibril structure and the resulting biological activities observed in our study.

## Accession numbers

PDB ID referenced in this manuscript : 4IMN

## Abbreviations

9,10-PQ: 9,10-Phenanthrenequinone;
AD: Alzheimer’s disease;
Aβ: Amyloid beta;
CPM: Chlorpheniramine;
L-PGDS: Lipocalin-type prostaglandin D synthase;
NMR: Nuclear Magnetic Resonance;
ThT: Thioflavin T;
TRD: Trazodone

## Acknowledgements

The research reported in this publication was supported by the Ministry of Education, AcRF Tier 2, Singapore under grant number M4020231.

The Electron microscopy work was undertaken at the NTU Institute of Structural Biology Cryo-EM lab in Nanyang Technological University, Singapore. We thank P. Padmanabhan and Y. Xia for their assistance in MTT cell assays.

