## Supplementary Figures for "Xenobiotic compounds modulate cytotoxicity of Aβ amyloids and interact with neuroprotective chaperone L-PGDS"

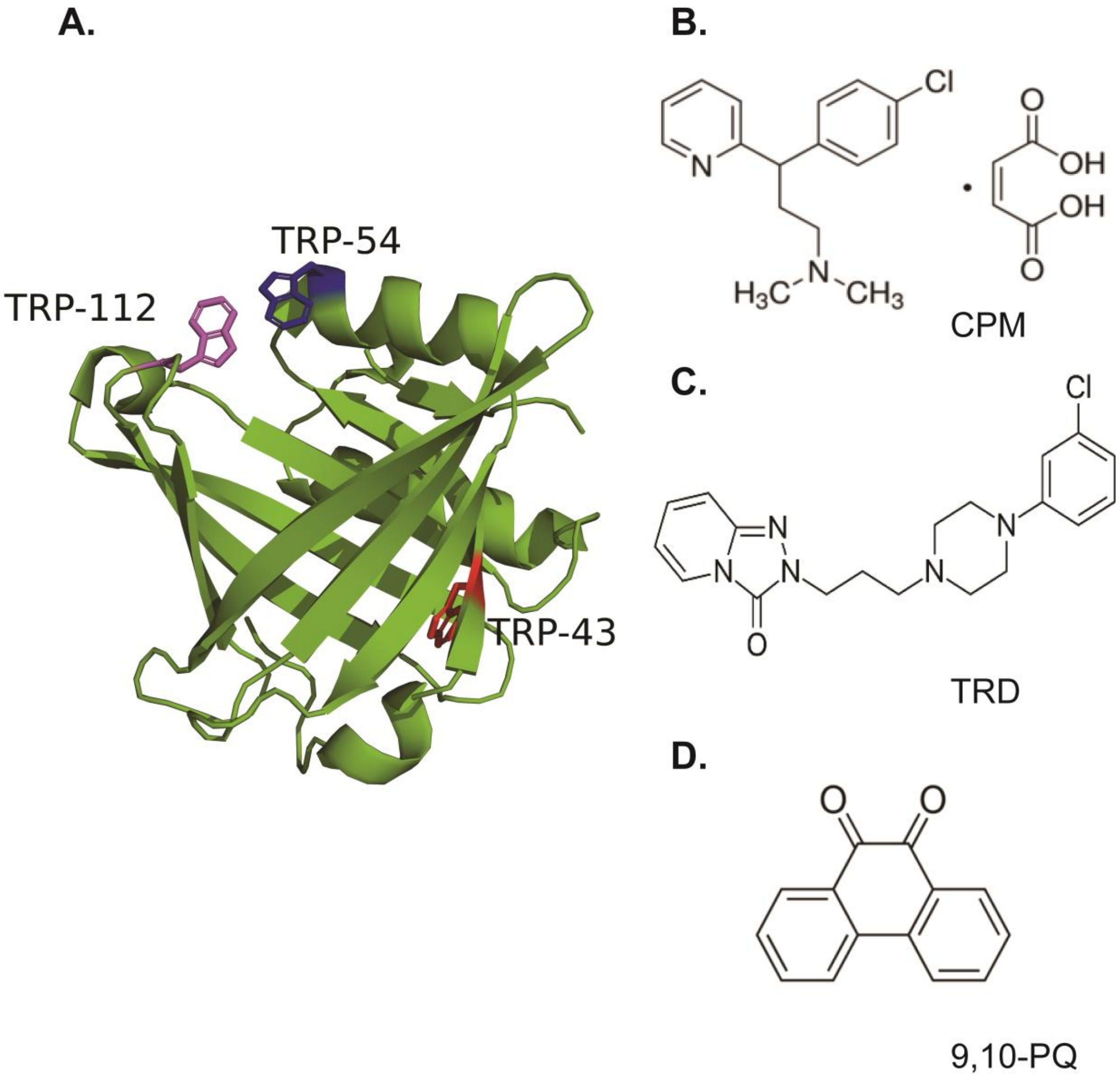


**Figure S1:** Structures of human WT L-PGDS and the anticholinergic drug compounds TRD and CPM. (A) Structure of human WT L-PGDS (PDB ID: 4IMN)[1]. The three different tryptophan residues of L-PGDS: $Trp_{43}$, $Trp_{54}$ and $Trp_{112}$ are shown in red, blue and magenta respectively. The chemical structures of CPM (B), TRD (C) and 9, 10-PQ (D).

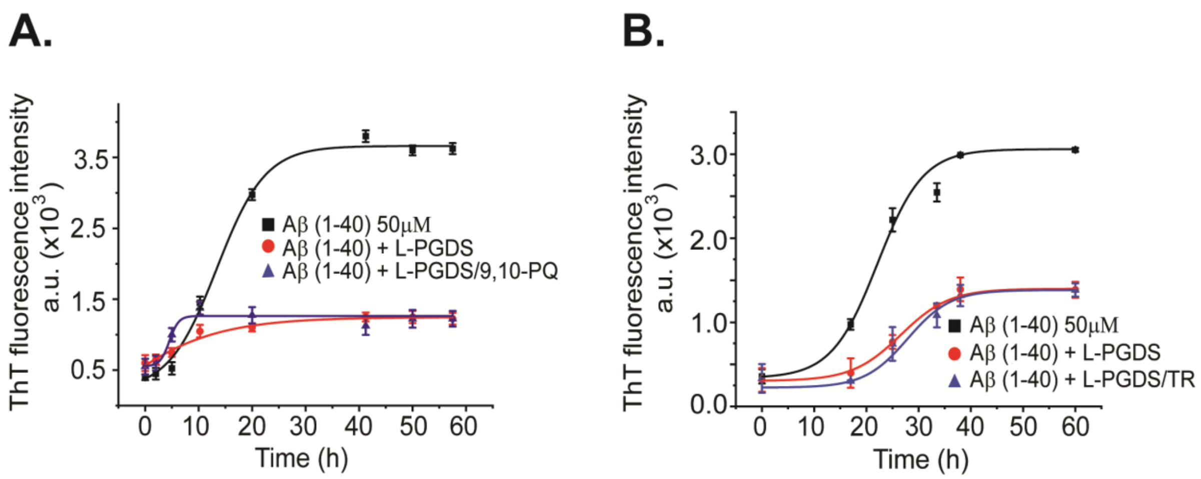


**Figure S2:** Inhibitory effects of L-PGDS and L-PGDS/drug complex on Aβ (1-40) peptide aggregation. (A) Representative time course of spontaneous Aβ (1-40) peptide aggregation in the absence (black) or presence of L-PGDS in 1:10 (protein: peptide) ratio (red) or presence of L-PGDS/9,10-PQ complex in ratio 1:1 (protein: compound) ratio (blue). (B) Representative time course of spontaneous Aβ (1-40) peptide aggregation in the absence (black) or presence of L-PGDS in 1:10 (protein: peptide) ratio (red) or presence of L-PGDS/TRD complex in 1:3 (protein: drug) ratio (blue).

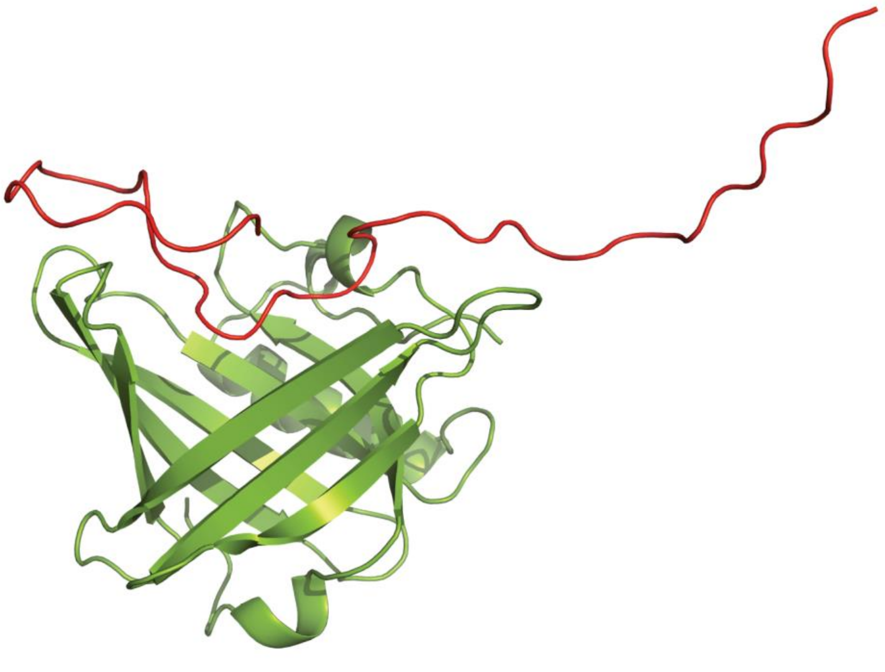

**Figure S3:** Figure reproduced from [2] 3D model of L-PGDS bound with Aβ obtained from molecular dynamics simulation. Structure of L-PGDS and Aβ (1-40) are represented in red and splitpea respectively.

[1] Lim SM, Chen D, Teo H, Roos A, Jansson AE, Nyman T, et al. Structural and dynamic insights into substrate binding and catalysis of human lipocalin prostaglandin D synthase. Journal of lipid research. 2013;54:1630-43.
[2] Kannaian B, Sharma B, Phillips M, Chowdhury A, Manimekalai MSS, Adav SS, et al. Abundant neuroprotective chaperone Lipocalin-type prostaglandin D synthase (L-PGDS) disassembles the Amyloid-β fibrils. Sci Rep. 2019;9.
